# Enhanced Solid Tumor Recognition and T cell Stemness with SynNotch CAR Circuits

**DOI:** 10.1101/2021.01.06.425642

**Authors:** Axel Hyrenius-Wittsten, Yang Su, Minhee Park, Julie M. Garcia, Nathaniel Perry, Garrett Montgomery, Bin Liu, Kole T. Roybal

## Abstract

The lack of highly tumor-specific antigens limits the development of engineered T cell therapeutics because of life-threatening “on-target/off-tumor” toxicities. Here we identify ALPPL2 as a tumor-specific antigen expressed in a spectrum of solid tumors, including mesothelioma. ALPPL2 can act as a sole target for chimeric antigen receptor (CAR) therapy or be combined with tumor-associated antigens such as MCAM or mesothelin in synthetic Notch (synNotch) CAR combinatorial antigen circuits. SynNotch CAR T cells display superior tumor control when compared to CAR T cells to the same antigens by prevention of CAR-mediated tonic signaling allowing T cells to maintain a long-lived memory and non-exhausted phenotype. Collectively, we establish ALPPL2 as a clinically viable target for multiple solid tumors and demonstrate the multi-faceted therapeutic benefits of synNotch CAR T cells.

**ONE SENTENCE SUMMARY:** SynNotch CAR circuits targeting novel solid tumor antigens enhance specificity and improve therapeutic efficacy by regulating T cell exhaustion.

## MAIN TEXT

Genetically modified T cells armed with chimeric antigen receptors (CARs) have shown unparalleled clinical efficacy in certain B cell malignancies by targeting lineage-restricted surface molecules (*1*). Translation of this success to solid tumors has hit roadblocks due to a lack of reliable tumor-specific antigens and sub-optimal therapeutic efficacy. One such treatment refractory solid tumor is mesothelioma — a highly aggressive chronic inflammation-driven cancer with an exceedingly poor prognosis (*2*). Mesothelioma is inherently hard to treat with traditional modes of cancer therapy and recent trials exploring mono- or combination immune checkpoint inhibitors (ICI) have also had limited impact (*3, 4*). CAR T cells targeting tumor-associated antigens such as mesothelin or fibroblast activation protein have shown preclinical promise, and are now in clinical trials alone or in combination with ICI (NCT02414269 and NCT01722149) (*5, 6*). However, we lack highly tumor-specific antigens with broad coverage of the three major mesothelioma subtypes: epithelioid (69% of cases, median overall survival ~14 months), biphasic (12%, ~10 months), and sarcomatoid (19%, ~4 months) (*7*).

We have now identified Alkaline Phosphatase Placental-like 2 (ALPPL2) as a highly specific and targetable cell surface antigen in all subtypes of mesothelioma (*8, 9*). ALPPL2 is not only relevant to mesothelioma but is also expressed on other solid tumor types. Immunohistochemistry (IHC) on tumor tissue arrays revealed that ALPPL2 is expressed in 43% of mesothelioma (39/91), 60% of ovarian cancer (36/60), 36% of pancreatic cancer (18/50), 18% of gastric cancer (13 of 72) and 100% of seminoma (11/11) (**Fig. 1A** and **Table S1**). Analysis of IHC data from the public protein expression database, the Human Protein Atlas, yielded similar results (**Table S1**).

**Figure 1.**
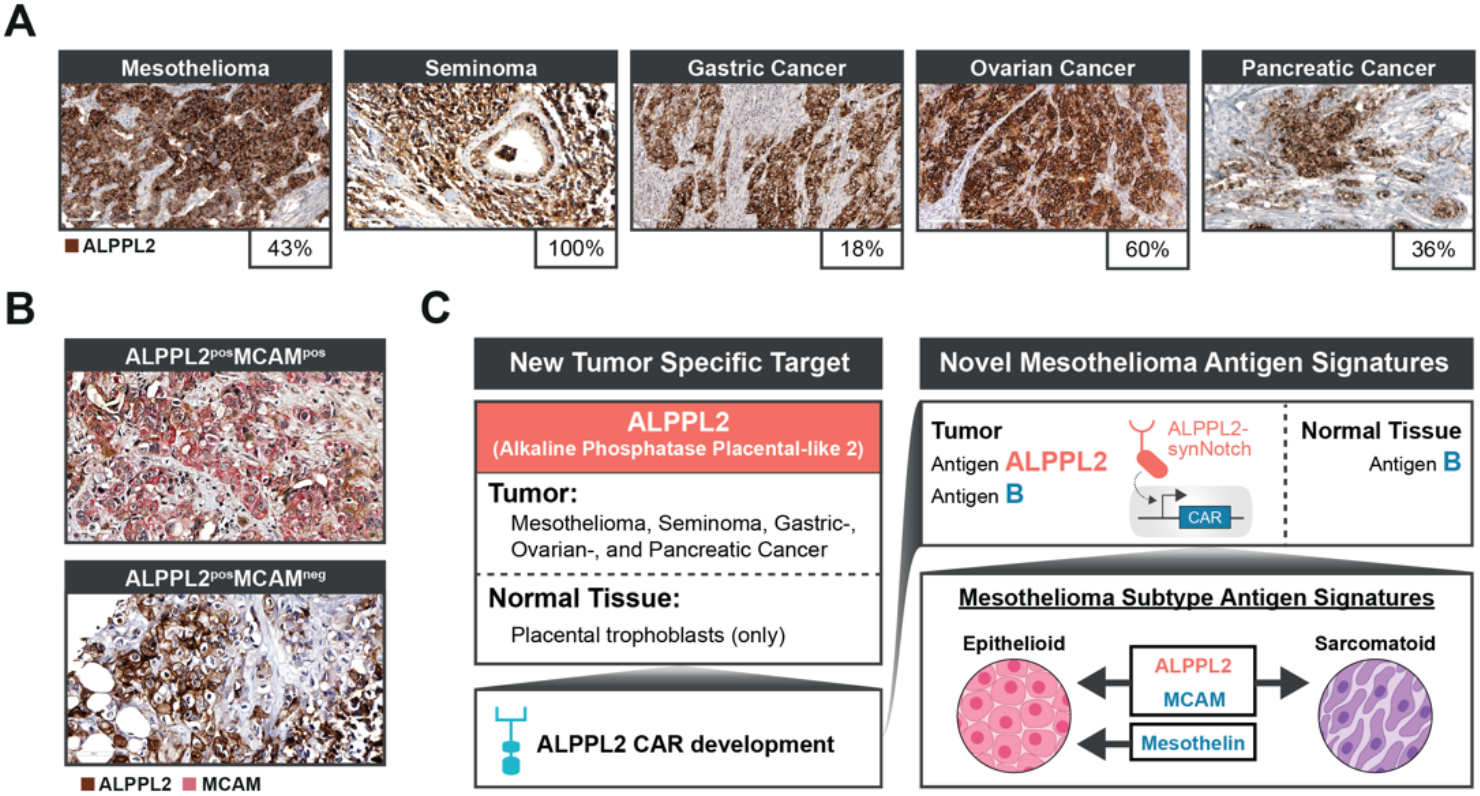
ALPPL2 - a highly specific and broadly applicable tumor antigen for combinatorial antigen recognition of solid tumors with SynNotch CAR T cells. (**A**) IHC study of ALPPL2 expression in FFPE solid tumor tissues. Positive staining was observed in mesothelioma, seminoma, gastric cancer, ovarian cancer, and pancreatic cancer. Scale bar: 100 μm for mesothelioma, pancreatic cancer, and gastric cancer; 200 μm for ovarian cancer and seminoma. (**B**) Representative images of ALPPL2 and MCAM co-expression in mesothelioma tissues. Scale bar: 50μm. (**C**) ALPPL2 is a highly tumor-specific antigen found in a range of solid tumor types that display minimal expression in normal tissues. ALPPL2 can be targeted directly using an ALPPL2 CAR but can also function as a priming switch for CARs targeting other tumor-associated antigen through synNotch CAR circuits to minimize “on-target/off-tumor” toxicity. We identified ALPPL2/MCAM as an antigen signature with coverage across mesothelioma subtypes. Mesothelin is mainly restricted to epithelioid mesothelioma.

To design a CAR targeting ALPPL2, we utilized an ALPPL2 binding single-chain variable fragment M25 (*8, 9*). Second generation anti-ALPPL2 M25 scFv 4-1BB-based (BBζ) CARs or two affinity matured clones (M25^ADLF^ and M25^FYIA^) (*9*) all effectively triggered activation of CD8^+^ T cells when stimulated with ALPPL2 positive tumor cells. With the exception of M25^ADLF^, CARs carrying M25 or M25^FYIA^ responded with high specificity to ALPPL2 and not the closely related homolog (89%) ALPI expressed in the healthy intestine (**Fig. S1A**). To further verify the tissue specificity of M25^FYIA^, an FDA-approved human tissue array that covers 20 different organs in duplicates (40 tissue cores total) was stained with M25^FYIA^ or a control non-binding scFv (YSC10). Positive staining by M25^FYIA^ was only observed in placental trophoblasts, with no staining observed in any other normal tissues (**Fig. S1B** and **Table S2**), in line with what we have previously observed for the corresponding full IgG1 antibodies (*9*). Although the functional role of ALPPL2 in tumor cells is unknown, ALPPL2 expression was recently described to be associated with and functionally essential for the establishment and maintenance of naïve pluripotency in various types of human pluripotent stem cells (*10*). The broad tumor expression and highly restricted normal tissue expression of ALPPL2, combined with the proposed function of ALPPL2 in various types of stem cells indicates that it might serve as an oncofetal antigen and carry high potential as immunotherapeutic target.

Advances in synthetic biology and immune cell engineering have led to approaches to engineer combinatorial antigen sensing capabilities into therapeutic T cells (*11*). We have previously engineered a new class of synthetic receptors based on the Notch receptor we call synNotch that enables custom gene regulation in response to a tissue or disease-related antigenic cue. SynNotch receptors can be engineered to sense a tumor antigen and induce the expression of a CAR to a second tumor-related antigen (*12*). These synNotch → CAR inducible circuits confine CAR expression and thus T cell activation to the site of disease. Here we sought to develop a clinically relevant synNotch → CAR circuit for mesothelioma that senses the combination of the novel tumor-specific antigen ALPPL2 and melanoma cell adhesion molecule (MCAM, also known as CD146 or MUC18) a mesothelioma-associated antigen found expressed in both epithelioid and sarcomatoid mesothelioma, as well as tumor associated blood vessels (*8, 13*). MCAM is reported to be restricted to few normal tissues (*14, 15*). To determine the co-expression pattern of ALPPL2 and MCAM, we performed dual-antibody IHC on human mesothelioma arrays. Using two different MCAM antibodies (**Table S3**), we observed ~52-81% MCAM co-staining in ALPPL2 positive tissue cores (**Fig. 1B** and **Table S4**). ALPPL2/MCAM cell surface co-expression was also seen on the mesothelioma cell lines M28 (epithelioid) and VAMT-1 (sarcomatoid) using our anti-ALPPL2 scFv (M25^FYIA^) and anti-MCAM scFv (M1) (**Fig. S1C**) (*16*). Indeed, ALPPL2 CAR expressing CD8^+^ T cells effectively recognized and killed both mesothelioma cell lines (**Fig. S1D**). Together, these results suggest that the majority of ALPPL2 expressing tumors also express MCAM, providing a rationale for dual targeting of ALPPL2/MCAM using an ALPPL2-synNotch design (**Fig. 1C**). Additionally, for patients with epithelioid mesothelioma, an alternative opportunity would be to use the well-established mesothelioma biomarker mesothelin as a secondary target, thereby reducing the risk of “on-target/off-tumor” toxicity associated with mesothelin (*17*). This approach would be mainly restricted, but widely applicable, to epithelioid mesothelioma for which mesothelin is found expressed in 84-93% of patients (*18, 19*) (**Fig. 1C**).

We first constructed BBζ CARs targeting either MCAM or mesothelin as well as a CD28 based (28ζ) mesothelin CAR, which is currently in clinical trials (**Fig. S2A** and **S2B**). CD8^+^ T cells engineered with the novel CARs and reference CD19 and mesothelin CARs were challenged with tumor cells expressing the corresponding antigens revealing specific activation, robust T cell proliferation, and production of Th_1_ immunostimulatory cytokines IL-2, IFN-γ, and TNFa (**Fig. S2C**, **S2D**, **S2E**, **S2F**, and **S2G**). To confine CAR expression locally to mesothelioma tumor tissues, we generated an ALPPL2-sensing synNotch using the M25^FYIA^ scFv and linked it to inducible genetic circuits containing either the MCAM CAR (BBζ) or a mesothelin CAR (BBζ or 28ζ) (**Fig. 2A**). CD8^+^ T cells equipped with these circuits were all able to selectively drive expression of the respective CARs when stimulated with target cells expressing ALPPL2 (**Fig. 2B**). As with T cells expressing the CARs constitutively, both the ALPPL2 synNotch → MCAM CAR and mesothelin CAR circuits induced robust T cell activation, proliferation, Th_1_ cytokine production, and tumor cell killing when combined with target cells expressing the correct ligand combination (**Fig. S3A**, **S3B**, **S3C,** and **S3D**).

**Figure 2.**
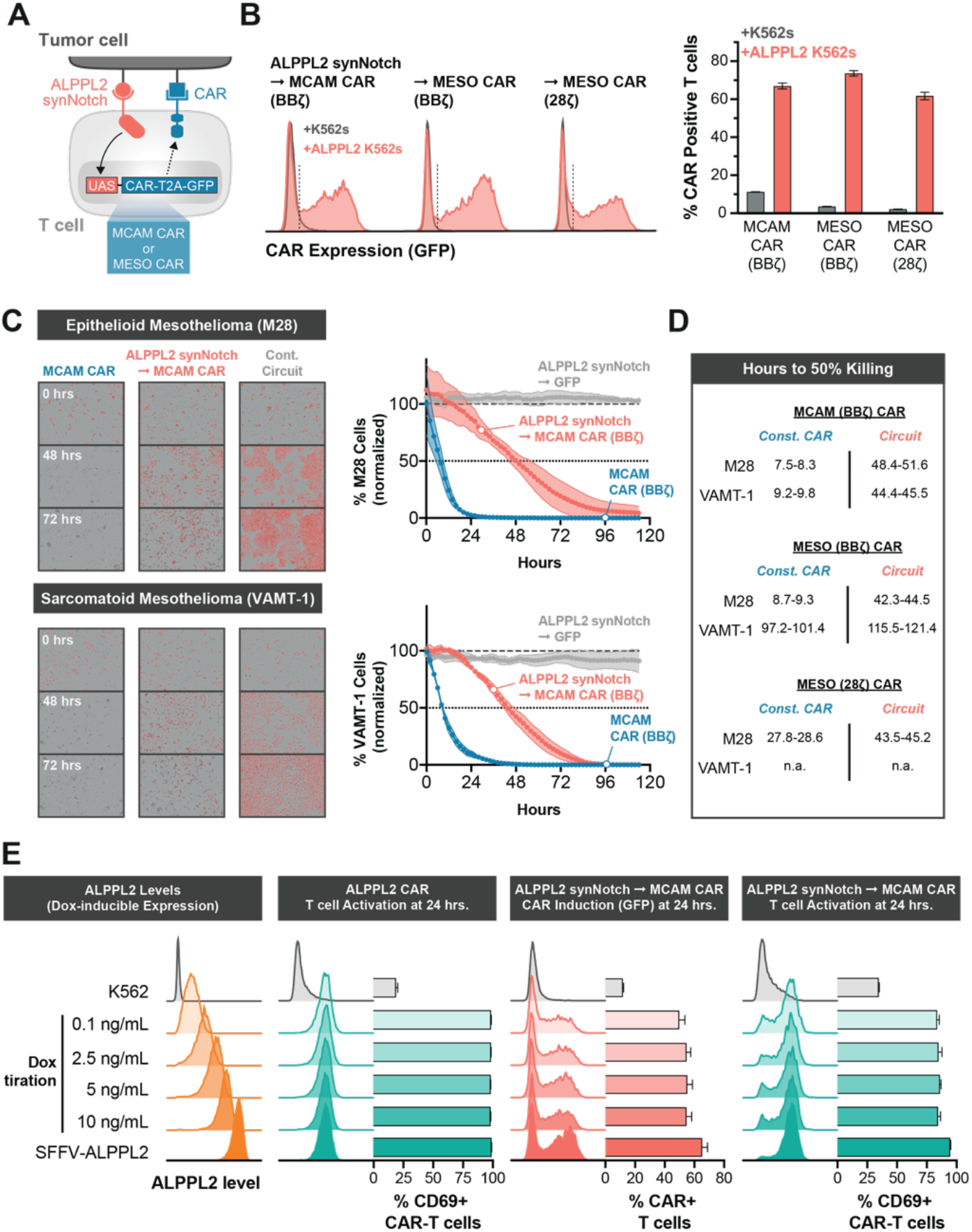
SynNotch CAR circuit T cells exhibit highly sensitive multi-antigen specificity and paced elimination of Mesothelioma. (**A**) Primary human CD8^+^ T cells engineered with ALPPL2 sensing synNotch with genetic circuits encoding for CARs targeting either MCAM (BBζ) or mesothelin (BBζ or 28ζ) CARs. (**B**) Antigen specific CAR expression was determined by GFP levels after 24hrs of stimulation with K562s expressing MCAM and mesothelin +/− ALPPL2 (**C**) Killing kinetics of epithelioid (M28) and sarcomatoid (VAMT-1) mesothelioma tumor cells by CD8^+^ T cells expressing a MCAM CAR constitutively or through an ALPPL2-synNotch circuit. Control circuit (Cont. circuit) is an ALPPL2-synNotch regulating GFP expression. (**D**) Hours to 50% killing of M28 and VAMT-1, as compared to untransduced CD8^+^ T cells, for constitutive or ALPPL2 synNotch regulated expression of MCAM (BBζ) or mesothelin (BBζ or 28ζ) CARs. (**E**) K562s constructed with doxycycline-inducible ALPPL2 expression showing dose-dependent induction of ALPPL2 after 72 hrs of doxycycline treatment. CD8^+^ T cells engineered with an ALPPL2 CAR or ALPPL2-synNotch MCAM CAR circuit were challenged with K562s displaying a range of ALPPL2 expression levels and analyzed for antigen specific CAR expression, as determined by GFP expression, and T cell activation, as determined by CD69 expression. n.a.; not applicable. Data is shown as mean±SD.

Both constitutive- and ALPPL2-primed expression of MCAM CAR was able to effectively elicit a cytotoxic response towards M28 and VAMT-1 tumor cell lines (**Fig. 2C**), whereas mesothelin CARs only targeted the epithelioid M28 cell line (**Fig. S3E**). There was a significant difference in the kinetics of cytotoxicity between constitutive- and circuit-controlled CAR expression, with the latter displaying several fold slower tumor killing rates (**Fig. 2D**). This more paced cytotoxic response of the synNotch → CAR circuit T cells can be attributed to the requirement for transcriptional regulation of CAR expression and as a population the circuit T cells do not upregulate the CAR in perfect unison. Protein levels of ALPPL2 vary within and between tumors based on IHC of various tumor types. Therefore, we wanted to confirm that our ALPPL2-synNotch remained responsive to lower antigen levels. For this purpose, we designed target cells for which levels of ALPPL2 could be controlled via titration of doxycycline (**Fig. 2E** and **S3F**). Using this system, we show that a low level of ALPPL2 is sufficient to induce robust expression of the MCAM CAR through the anti-ALPPL2 M25^FYIA^-based synNotch, allowing for potent T cell activation through combinatorial antigen sensing (**Fig. 2E**).

Another limiting factor in CAR T therapy of solid tumors is the intrinsic maintenance of multifunctional T cell states and the prevention of T cell exhaustion/dysfunction. In the absence of antigen, constitutive expression of CARs is known to elicit tonic signaling. This low level signaling is linked to detrimental effects on the phenotypic state of CAR T cells, such as differentiation of long-lived T cell memory phenotypes towards short-lived effector states and the upregulation of inhibitory receptors (*20–22*). Although most of the components of CAR domain architecture impact tonic signaling (e.g. the scFv, hinge, and signaling domains), surface expression levels remain as the unifying determinant (*20*). We thus reasoned that restricting the timing of CAR expression and location of expression to the tumor tissue with synNotch could prevent CAR-mediated tonic signaling and the detrimental effects thereof. We observed that constitutive expression of the MCAM CAR in CD8^+^ T cells impacted T cell differentiation and consistently displayed a significantly smaller fraction of long-lived T stem cell memory (T_SCM_) cells (defined as CCR7^+^CD45RO^-^CD27^+^CD45RA^+^CD95^+^) and a higher proportion of T effector memory cells (defined as CCR7^-^CD45RO^+^) in contrast to the ALPPL2 synNotch → MCAM CAR circuit (**Fig. 3A** and **3B**). The same protection from premature T cell differentiation was observed when comparing ALPPL2-synNotch → ALPPL2 CAR or CD19 CAR circuits T cells to T cells that constitutively express the corresponding CARs. Thus, synNotch-mediated CAR regulation is a general means to control tonic signaling and prevent detrimental T cell differentiation prior to antigen exposure, and this effect remains during long-term *ex vivo* T cell culture. (**Fig. 3C** and **S4A**). In line with this, constitutive CAR expression led to higher expression and co-expression of surface markers linked to T cell exhaustion that was avoided by synNotch transcriptional regulation (**Fig. 3D**, **S4B**, **S4C**, and **S4D**). Upon target cell stimulation, MCAM CAR expression levels were comparable between constitutive and ALPPL2 synNotch transcriptional regulation (**Fig. S4E**). Further, Jurkat T cell reporter systems for AP-1, NFAT, and NF-κB transcriptional activity showed that constitutive expression of the MCAM CAR was sufficient to induce NF-κB-mediated transcription (**Fig. 3E** and **S4F**). NF-κB transcriptional activity was not observed when the MCAM CAR was under control of the ALPPL2 synNotch suggesting that synNotch regulated CAR expression maintains the engineered T cell population in a superior functional state by circumventing unfavorable tonic signaling. NF-κB signaling has previously been associated with tonic signaling in BBζ CARs and has been linked to CAR expression levels (*23*).

**Figure 3.**
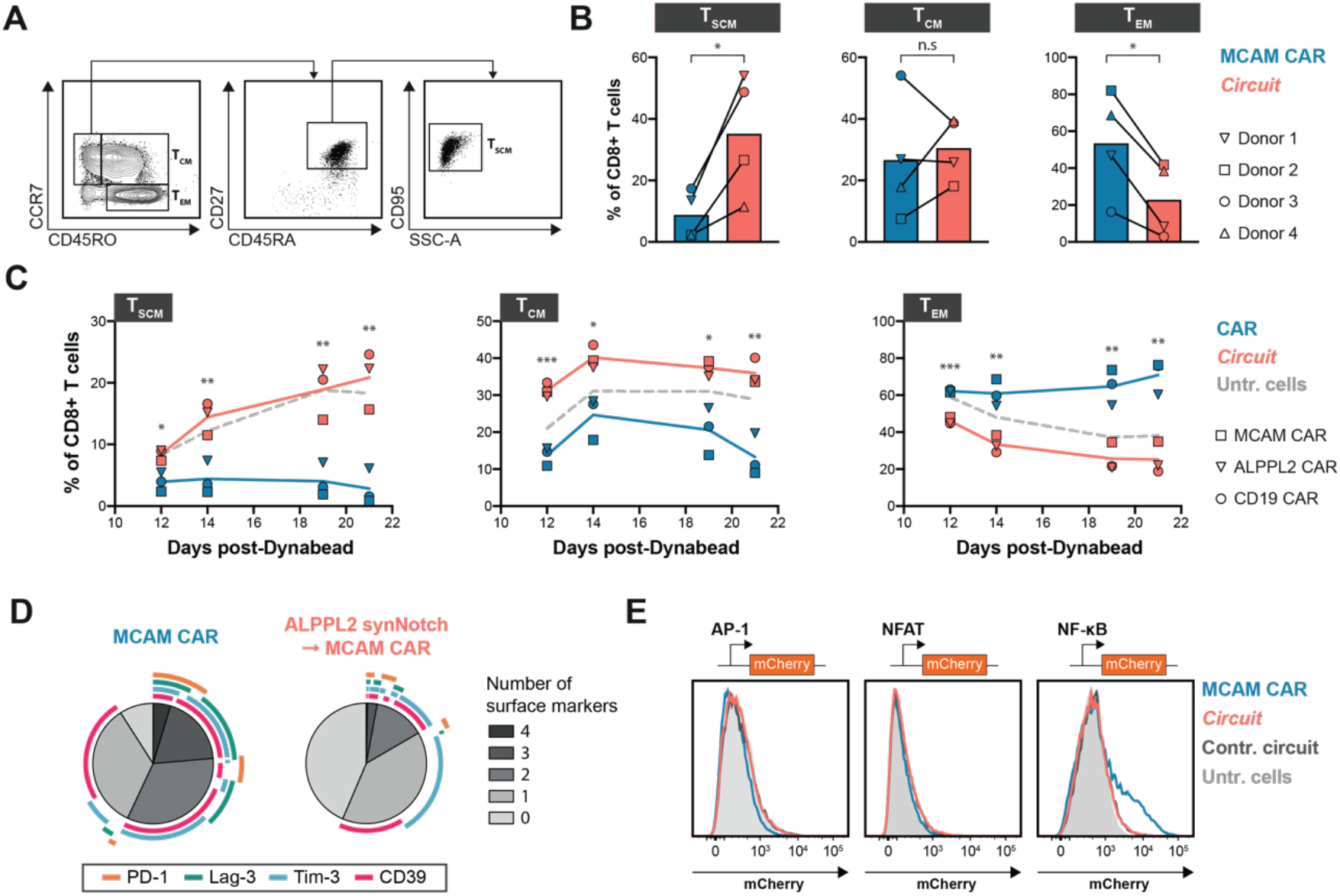
SynNotch regulation of CAR expression is a general means to maintain T cell stemness prior to therapeutic administration. (**A**) Gating strategy for identifying T stem cell memory (T_SCM_), T central memory (T_CM_), and T effector memory (T_EM_) cells. (**B**) Composition of T_SCM_, T_CM_, and T_EM_ in non-antigen exposed CD8^+^ T cells from four different donors engineered to express a MCAM CAR either constitutively or through an ALPPL2-synNotch circuit 14 days post initial CD3/CD28 Dynabead stimulation. (**C**) Longitudinal T cell memory subsetting of donor 4 engineered to express either an MCAM, ALPPL2, or CD19 CAR constitutively or through ALPPL2-synNotch circuits. (**D**) Expressional pattern analysis of CD39, Lag-3, PD-1, Tim-3 in non-antigen exposed CD8^+^ T cells from three different donors engineered to express a MCAM-CAR either constitutively or through ALPPL2 synNotch circuit 14 days post initial CD3/CD28 Dynabead stimulation. (**E**) Jurkats carrying AP-1, NFAT, or NF-kB response elements expressing a MCAM CAR constitutively or through ALPPL2-synNotch circuit. Control circuit is ALPPL2 synNotch driving GFP expression. Statistics was calculated using paired (**B**) or unpaired (**C**) Student’s t-test. *P ≤0.05; **P ≤0.01; ***P ≤0.001; n.s.; not significant.

To evaluate the therapeutic efficacy of our new CARs and clinically viable synNotch circuits, NSG mice were implanted with epithelioid M28 tumors prior to receiving an infusion of ALPPL2 CAR T cells, MCAM CAR T cells, or ALPPL2 synNotch → MCAM-CAR circuit T cells. Constitutive MCAM CAR T cells exhibited inconsistent ability to control M28 tumor growth, whereas ALPPL2 CAR T showed significant reduction in tumor growth (**Fig. 4A**). However, the greatest effect was observed for the ALPPL2 synNotch → MCAM CAR circuit T cells, for which a complete response was observed in a majority of the mice (**Fig. 4A**).

**Figure 4.**
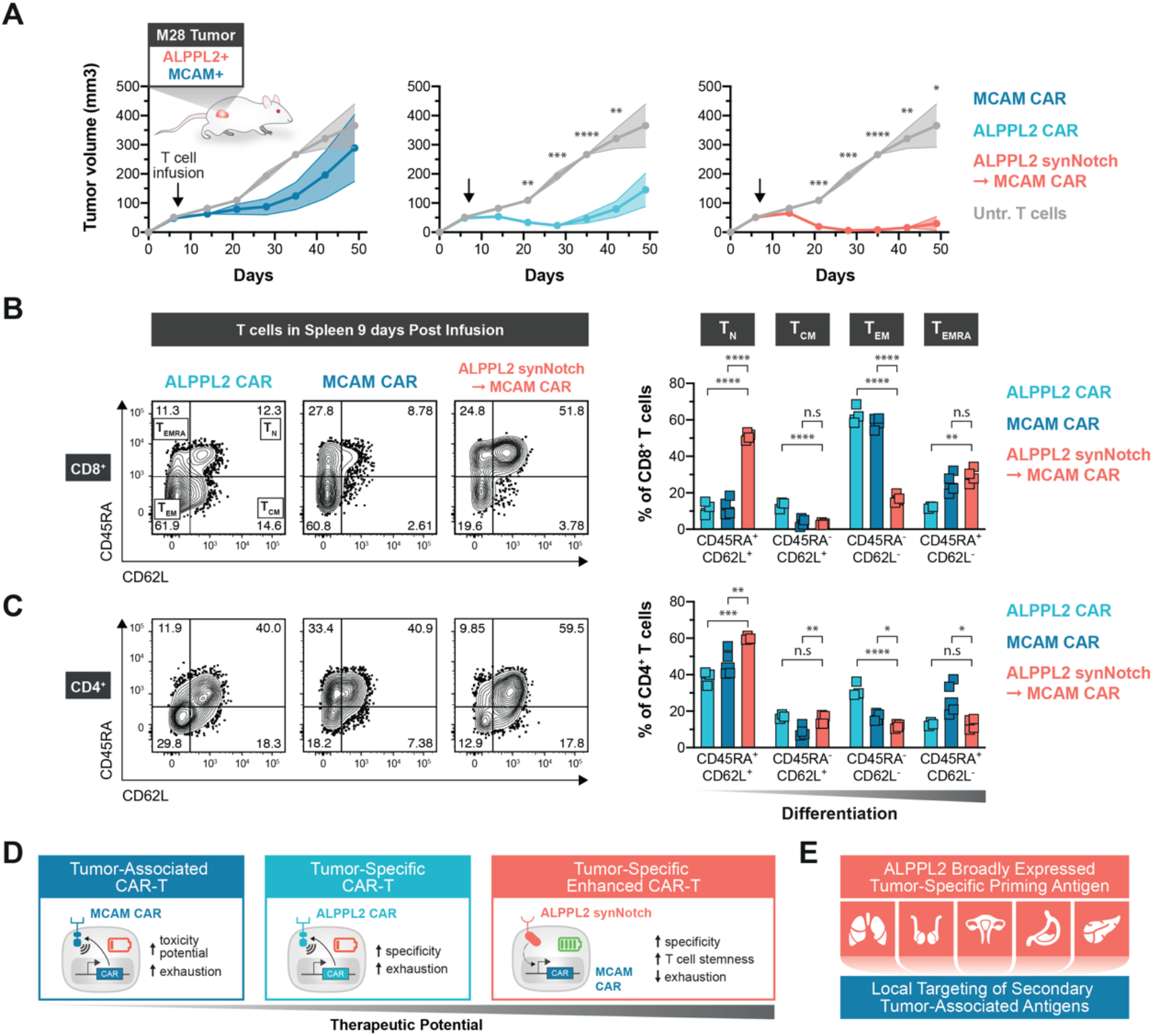
SynNotch CAR Circuit T cells exhibit superior efficacy and persistence in vivo. (**A**) NSG subcutaneously implanted with M28 tumor cells were injected i.v. with 1:1 ratio of CD4^+^:CD8^+^ T cells engineered with an ALPPL2 CAR (n=5), MCAM CAR (n=5), ALPPL2 synNotch regulating MCAM-CAR expression, or untransduced T cells (n=5). Tumor size was monitored over 49 days. Data is presented as mean±SEM. (**B**,**C**) Percentage of T cell memory subsets in (**B**) CD8^+^ and (**C**) CD4^+^ in the spleen 9 days after T cell infusion. (**D**) SynNotch CAR circuits allow for safe targeting of tumor-associated antigens and enhances immunotherapeutic potential by avoiding tonic CAR signaling outside tumor tissues. (**E**) ALPPL2 is a novel highly tumor-specific antigen expressed in mesothelioma, seminoma, gastric-, ovarian-, and pancreatic cancer that allows for safe targeting of other less specific tumor-associated antigens. Statistics were calculated using either two-way ANOVA with Tukey’s multiple comparison (**A**) or one-way ANOVA with Dunnett’s multiple comparisons test (**B**,**C**). *P ≤0.05; **P ≤0.01; ***P ≤0.001; ***P <0.0001; n.s.; not significant.

We then sought to determine the underlying reasons for the enhanced efficacy of the synNotch → CAR circuit T cells. We assessed CD4+ and CD8+ engineered T cells in the spleen and determined that the number of T cells correlated with the ability to control tumor growth (**Fig. S5A**), suggesting that T cell persistence plays a pivotal role in therapeutic efficacy. T cell persistence inversely correlated with the level of tonic signaling (fraction CD39+) in the CD4^+^ and CD8^+^ pre-infusion ALPPL2 synNotch circuit and constitutive CAR T cells (MCAM CAR > ALPPL2 CAR > circuit, **Fig. S5B**). In line with this, ALPPL2 → MCAM CAR circuit T cells harvested 9 days post infusion from the spleen were more enriched for long-lived memory phenotypes in both the CD8^+^ and CD4^+^ subsets compared constitutive CAR T cells (**Fig. 4B** and **4C**).

Here we have explored the multidimensional advantages of synNotch → CAR circuits, which circumvent some of the critical problems thought to prevent efficacy of cell therapies for solid tumors. SynNotch → CAR circuits not only provide improved specificity through multi-antigen sensing, but are also a general means to enhance therapeutic efficacy through cell-autonomous and context-dependent regulation of CAR expression and prevention of tonic signaling, leading to the maintenance of T cell memory subsets important for persistence and sustained activity of the cell therapy (**Fig. 4D**). We also identify ALPPL2 as a highly promising tumor-specific antigen for engineered T cell therapies for a range of solid tumors. ALPPL2 can act as a singular target for CAR T cell therapy or can be sensed in combination with secondary highly expressed tumor-associated antigens via synNotch → CAR circuits to maximize tumor elimination and reduce toxicity potential (**Fig. 4E**).

## NOTES

## Acknowledgements

A.H-W. has received funding from the Swedish Society for Medical Research and the Swedish Research Council. K.T.R. is funded by the Parker Institute for Cancer Immunotherapy, the UCSF Helen Diller Family Comprehensive Cancer Center, the Chan Zuckerberg Biohub, an NIH Director’s New Innovator Award (DP2 CA239143), Cancer Research UK, and the Kleberg Foundation. We acknowledge the PFCC supported in part by Grant NIH P30 DK063720 and by the NIH S10 Instrumentation Grant S10 1S10OD021822-01.

## Author Contributions

A.H.W. designed the study, designed and performed experiments and vector construction, analyzed data, and wrote the manuscript; Y.S. performed and analyzed histological stains, assisted in *in vivo* experiments, and wrote the manuscript; M.P., N.P., and G.M. performed vector construction; M.P. and J.G. performed *in vitro* experiments; B.L. and K.T.R. conceived and designed the study, designed experiments, and wrote the manuscript.

## SUPPLEMENTARY MATERIALS

Materials and Methods

Tables S1–S4

Figs. S1–S4

## SUPPLEMENTARY MATERIALS

### MATERIALS AND METHODS

#### Recombinant scFv production

ScFv expression and purification was done as previously described (*24, 25*). Briefly, the scFv gene was cloned into the secretion vector pUC119mycHis to impart a c-myc epitope and a hexahistidine tag at the C-terminus. Soluble scFv was harvested from the bacterial periplasmic space and purified by immobilized metal affinity (HiTrap His, GE HealthCare) and ion-exchange (DEAE, GE HealthCare) chromatography.

#### Immunohistochemistry

IHC studies on frozen tissues were performed as described (*9*). Briefly, frozen normal human tissue arrays (US Biomax) were air-dried for 15 min at RT, fixed in 4% paraformaldehyde for 10 min, washed 3 times with PBST (PBS containing 0.1% Tween-20), and incubated with 3% H_2_O_2_ (Thermo Fisher Scientific) for 10 min to block endogenous peroxidase activity, washed with PBST, further blocked with PBST containing 2% goat serum (Jackson ImmunoResearch Laboratories) and 5% BSA (Thermo Fischer Scientific) at RT for 1h. After avidin/biotin (Vector Laboratories) blocking and washed three times with PBST, slides were incubated with 10 μg/ml biotinylated scFvs (M25^FYIA^ and a non-binding C10) at 4 °C overnight, followed by detection with streptavidin-HRP (Jackson ImmunoResearch Laboratories) using DAKO liquid DAB+ (Agilent). Slides were counterstained with hematoxylin followed by bluing reagent (Scytek), mounted in Aqua-mount (Lerner Laboratories), and scanned by Aperio AT2 digital scanner (Leica).

IHC study on FFPE tissue arrays were performed as described (*9*). Briefly, FFPE slides were deparaffinized in xylene overnight, rehydrated by sequential exposure to 100%, 95%, 70% ethanol, and ddH_2_O (7 min each), and placed in Tris Buffer (10 mM Tris Base, 1 mM EDTA, 0.05% Tween 20, pH 9.0) at 95°C for 25 min for antigen retrieval, washed three times with PBST, sequential blocked with 3% H_2_O_2_ and 2% donkey serum (Santa Cruz Biotechnology Inc), incubated at 4°C overnight with primary antibodies (anti-ALPPL2 mouse mAb (LifeSpan clone SPM593, catalogue# LS-C390148-20) at 0.4 μg/ml, or anti-MCAM rabbit mAb (Abcam clone EPR3208, catalogue# ab75769) at 0.625 μg/ml, or anti-MCAM rabbit polyclonal antibody (Abcam, catalogue# ab228487) at 10 μg/ml), washed three times with PBST, further incubated with anti-mouse DAKO EnVision+ (Agilent) or anti-rabbit DAKO EnVision+ (Agilent), followed detection using DAKO liquid DAB+ (Agilent). Slides were counterstained with hematoxylin followed by bluing reagent (Scytek), dehydrated by sequential exposure to 70%, 95%, 100% ethanol, mounted using Permount mounting media (Thermo Fischer Scientific) and scanned by Aperio XT and AT2 digital scanners (Leica).

For MCAM/ALPPL2 co-staining on FFPE tissue arrays (US Biomax), the ImmPRESS^®^ Duet Double Staining HRP/AP Polymer Kit was used (HRP anti-Mouse IgG and AP anti-Rabbit IgG, Vector Laboratories). Following deparaffinization and antigen retrieval with Tris Buffer (10 mM Tris Base, 0.05% Tween 20, pH 10.0), endogenous peroxidase activity was blocked by BLOXALL blocking solution for 15 min followed by PBST wash. The slides were further blocked with 2.5% normal horse serum at RT for 1h, washed three times with PBST and incubated with the following primary antibody pairs diluted into 2.5% normal horse serum and incubated with the tissue slides at 4°C overnight: pair 1: anti-MCAM rabbit mAb and anti-ALPPL2 mouse mAb (both at 1:200 dilution); and pair 2: anti-MCAM rabbit pAb and anti-ALPPL2 mouse mAb (both at 1:100 dilution). Slides were washed three times with PBST, incubated with ImmPRESS Duet reagent at RT for 30 min, followed by sequential detection with ImmPACT DAB EqV and ImmPACT Vector Red substrates. Slides were counterstained with hematoxylin followed by bluing reagent (Scytek). After dehydration, the slides were mounted using Permount mounting media (Thermo Fischer Scientific) and scanned by Aperio XT and AT2 digital scanners (Leica).

#### Vector Construction Designs

All CARs contained CD8α signaling peptide (MALPVTALLLPLALLLHAARP) followed by a FLAG tag (DYKDDDDK) and either an anti-CD19 (FMC63) (*26*), anti-ALPPL2 (M25, M25^ADLF^, or M25^FYIA^) (*9*), anti-MCAM (M1 or M40) (*13*), or anti-mesothelin (m912) (*27*) scFv. CARs were then fused to either the CD8α hinge/transmembrane domain, 4-1BB intracellular domain, and CD3ζ intracellular domain or CD28 hinge/transmembrane domain and CD3ζ intracellular domain. In order to determine expression, a T2A self-cleaving peptide preceding eGFP was attached to CAR coding sequence. In some experiments, CARs with a c-Myc-tag (EQKLISEEDL) instead of the FLAG tag and a GS linker tethered eGFP instead of T2A-eGFP were used. All constitutive CARs were cloned into a modified pHR’SIN:CSW vector containing a SFFV promoter. The ALPPL2 synNotch was constructed as previously described (*12*) using the anti-ALPPL2 (M25^FYIA^) scFv. The ALPPL2-synNotch was cloned into a modified pHR’SIN:CSW vector containing a PGK promoter. Inducible CAR response elements were constructed by subcloning the CAR-T2A-eGFP sequences into a modified pHR’SIN:CSW vector carrying five copies of the Gal4 DNA binding domain target sequence (GGAGCACTGTCCTCCGAACG) 5’ to a minimal CMV promoter. These vectors also included a PGK promoter constitutively driving the expression of either of the fluorescent reporters mCherry or BFP. For ectopic expression of ALPPL2, ALPI, and mesothelin, coding sequences corresponding to NP_112603.2 (ALPPL2), NP_001622.2 (ALPI), and NP_001170826.1 (mesothelin) were cloned into a modified pHR’SIN:CSW vector containing a SFFV promoter. For ALPPL2 and ALPI, a FLAG tag was inserted 3’ of their predicted signaling peptides (MQGPWVLLLLGLRLQLSLG for both). To establish non-perturbing nuclear fluorescent labeling, Sv40 NLS (PKKKRKV) was fused 5’ to mKate2 via a linker sequence (DPPVAT) and cloned into a modified pHR’SIN:CSW vector containing a EF1α promoter. To establish doxycycline controlled expression of ALPPL2, FLAG tagged ALPPL2 was cloned into a modified pHR’SIN:CSW vector containing a TRE3GS inducible promoter with constitutive downstream cassette of a SFFV promoter driving the expression of rtTA3. To enable genetic knockout (KO) of MCAM, the gRNA sequence GAGGCGCAGCTCCCGGGCTGG was cloned into the pL-CRISPR.EFS.GFP vector, which was a gift from Benjamin Ebert (Addgene plasmid #57818). AP-1, NFAT, and NFkB response elements and the minimal promoters used for T cell activity reporters were a gift from Peter Steinberger (Addgene plasmid #118031, 118094, and 118095) and were cloned together with mCherry into a modified pHR’SIN:CSW vector. All constructs were cloned via In-Fusion cloning (Takara Bio).

#### Lentiviral Production and Cell Lines

To produce pantropic VSV-G pseudotyped lentivirus, Lenti-X 293T cells (Takara Bio) were transfected with a pHR’SIN:CSW transgene expression vector and the viral packaging plasmids pCMVdR8.91 and pMD2.G using TransIT^®^-Lenti Transfection Reagent (Mirus Bio LLC). Lentivirus containing medium was harvested 48 hrs post-transfection. ALPPL2^+^, ALPI^+^, CD19^+^, and Mesothelin^+^ K562 (ATCC) target cells lines were generated by lentiviral transduction followed by fluorescence-activated cell sorting (FACS) based on cell surface protein expression. ALPPL2^+^ K562 cells were sorted based on FLAG tag expression. Generation of MCAM KO K562 target cells was performed by lentiviral transduction with a vector carrying sgRNA-Cas9-P2A-eGFP followed by FACS based on eGFP^+^CD146(MCAM)^-^ cells. Human mesothelioma cell lines, M28 (epithelioid) and VAMT-1 (sarcomatoid), were originally obtained from Dr. Brenda Gerwin’s lab at the National Cancer Institute. Mesothelin^+^ M28 target cells were generated by lentiviral transduction followed by FACS based on cell surface mesothelin expression. To enable real-time counting of M28 and VAMT-1 target cells, these cell lines were tagged with a non-perturbing fluorescent mKate2 reporter through lentiviral transduction followed FACS purification of mKate2+ cells. Inducible ALPPL2^+^ K562 target cells were generated via lentiviral transduction followed by single-cell sorting of doxycycline treated cells into 96-well U-bottom plates. Clonal lines were assessed for dose dependent ALPPL2 expression, as defined by cell surface FLAG tag levels, and the most uniform line was used for subsequent assays. T cell activation reporter lines for AP-1, NFAT, and NFkB were generated through lentiviral transduction of Jurkat (clone E6-1) cells (ATCC) and sorted based on the low basal activity seen for these reporters. Lenti-X 293T cells and M28 were cultured in DMEM (Gibco) with 10% fetal bovine serum (MilliporeSigma), 50units/mL Penicillin and 50 μg/ml Streptomycin (MP biochemicals), and 1X Sodium Pyruvate (MilliporeSigma). K562s were cultured in IMDM (Gibco) with 10% FBS, 50 units/mL Penicillin and 50 μg/ml Streptomycin. Jurkat, M28, and VAMT-1 were cultured in RPMI-1640 (Gibco) with 10% FBS, 100 units/mL Penicillin and 100 μg/ml Streptomycin, 1X Glutamax (Gibco).

#### Isolation and Engineering of Human T cells

Primary CD4^+^ and CD8^+^ T cells were isolated from Fresh Human Peripheral Blood Leukapheresis Packs (STEMCELL Technologies) using EasySep™ isolation kits (STEMCELL Technologies). T cells were cryopreserved in RPMI-1640 with 20% human AB serum (Valley Biomedical) and 10% DMSO (Fisher Scientific). After thawing, T cells were cultured in human T cell medium consisting of X-VIVO 15 (Lonza), 5% Human AB serum, and 10 mM neutralized N-acetyl L-Cysteine (MilliporeSigma) supplemented with 30 units/mL recombinant human IL-2 (R&D Systems). Primary human CD4^+^ or CD8^+^ T cells were thawed and left to recover for 24 hrs after which they were stimulated with Human T-Activator CD3/CD28 Dynabeads (Thermo Fisher Scientific) at a 1:1 cell:bead ratio. After an additional 24 hrs, the primary T cells were exposed to lentivirus for 24 hrs. At day 5 after T cell stimulation, the Dynabeads were removed and T cells were sorted to carry to correct transgene composition. T cells were expanded and rested until day 10 for *in vivo* experiments and day 14 for *in vitro* assays.

#### Doxycycline inducible ALPPL2

A clonal line of K562 cells carrying a doxycycline inducible ALPPL2 cassette was treated with doxycycline (Abcam) at doses ranging between 0.1-10 ng/mL for 3 days. Surface expression levels were assessed by flow cytometry through the FLAG-tag on ALPPL2 before being used in T cell stimulation assays.

#### Activation, Proliferation, and Intracellular Cytokine Assays

For *in vitro* stimulation assays of engineered T cells, effector cells were combined with targets cells at a 1:1 ratio in 96-well U-bottom plates and centrifuged for 1 min at 400 × g to force interaction of the cells. To assess T cell activation, cells were analyzed for surface expression of CD69 16-24 hrs post target challenge. For proliferation assays, engineered T cells were stained with CellTrace Violet (CTV) Cell Proliferation Kit (Thermo Fisher Scientific) prior to being combined with target cells. CTV dilution was assessed 4 days post target challenge. Intracellular cytokine (IC) assays using constitutive CARs, were set up as above but in the presence of 1X Brefeldin A solution (Thermo Fisher Scientific) and 1X Monensin solution (Biolegend). IC assays using inducible CAR circuits were performed by incubating effector cells and targets for 16 hrs before the addition of 1X Brefeldin A solution and 1X Monensin solution, in order to enable robust CAR expression and trafficking to the cell surface before blocking protein transport. All experiments were performed in T cell medium supplemented with 30 units/mL IL-2.

#### Incucyte Killing Assay

For *in vitro* engineered T cell killing assays, M28 or VAMT-1 expressing nuclear mKate2 were seeded in 96-well flat-bottom plates. After 24 hrs engineered T cells were added at an expected effector:target ratio of 2:1. Plates were imaged every 2 hrs using the IncuCyte^®^ S3 Live-Cell Analysis System (Essen Bioscience) for a duration of 5 days. Three images per well at 10X magnification were collected and analyzed using the IncuCyte^®^ S3 Software (Essen Bioscience) to detect and count the number of mKate2^+^ nuclei per image. Experiments were performed in RPMI-1640 with 10% FBS, 100 units/mL Penicillin and 100 μg/ml Streptomycin, 1X Glutamax supplemented with 30 units/mL IL-2.

#### Flow Cytometry Antibodies, Analysis, and Sorting

For scFv binding analysis, M28 and VAMT-1 mesothelioma cells in exponential growth phase were incubated with M1 (MCAM) or M25^FYIA^ (ALPPL2) scFv for 1h at RT, washed three times with PBS, further incubated with a secondary anti-6xHis (4E3D10H2/E3, MA1-135-A647, Thermo Fischer Scientific) at 1:1000 dilution for 1h at RT, and washed three times with PBS before analysis. For immunophenotyping, samples were stained for either 20-30min at 4°C, 30min at RT, or 15min at 37°C followed by 15min at RT, depending on the panel. For extracellular stains, cell washes and final resuspension was performed with PBS supplemented with 2% FBS. IC stains were performed using an Intracellular Fixation & Permeabilization Buffer Set (Thermo Fisher Scientific) per manufacturer’s instructions. Dead cells were excluded with Draq7 (Abcam) or Zombie NIR™ Fixable Viability Kit (Biolegend).

For immunophenotyping, the following antibodies were used: anti-CD146 (541-10B2, 130-097-942, Miltenyi), anti-CD19 (HIB19,11-0199-41, eBioscience), anti-CD197 (G043H7, 353226, Biolegend), anti-CD223 (11C3C65, 369322, Biolegend), anti-CD27 (M-T271, 356424, Biolegend), anti-CD279 (EH12.2H7, 329906 and 329908, Biolegend), anti-CD3 (UCHT1, 300463, Biolegend), anti-CD366 (7D3, 563422, BD biosciences), anti-CD366 (F38-2E2, 345026, Biolegend), anti-CD39 (A1, 328228, Biolegend), anti-CD4 (SK3, 563875, BD biosciences), anti-CD45 (2D1, 368515, Biolegend), anti-CD45RA (HI100, 304150, Biolegend), anti-CD45RO (UCHL1, 564291, BD biosciences), anti-CD62L (DREG-56, 304830 and 304822, Biolegend), anti-CD69 (FN50, 564364, BD biosciences), anti-CD69 (FN50, 310910, Biolegend), anti-CD8a (RPA-T8, 563796 and 563823, BD biosciences), anti-CD8a (SK1, 344706, Biolegend), anti-CD8a (OKT8, 17-0086-42, eBioscience), anti-CD95 (DX2, 305644, Biolegend), anti-FLAG (DYKDDDDK) tag (L5, 637310, Biolegend), anti-IFN-*γ* (4S.B3, 563731, BD biosciences), anti-IL-2 (MQ1-17H12, 560707, BD biosciences), anti-Mesothelin (REA1057, 130-118-096, Miltenyi), anti-Myc-tag (9B11, 2233S, Cell Signaling Technology), and anti-TNF (MAb11, 563996, BD biosciences). Cells were analyzed using either Accuri™C6, LSR II SORP, FACSymphony X50 SORP or, for cell sorting, FACSAria II SORP, FACSAria IIIu SORP, or FACSAria Fusion SORP (all BD Biosciences). Cell counts were performed using flow cytometry with CountBright Absolute Counting Beads (Thermo Fisher Scientific) per manufacturer’s instructions. Data was analyzed using FlowJo software (BD Biosciences).

#### Xenograft Tumor Models

NOD.Cg-*Prkdc^scid^Il2rg^tm1Wjl^*/SzJ (NSG) mice were implanted with 4 x 10^6^ M28 tumor cells subcutaneously of the right flank. Seven days after tumor implantation, 3 x 10^6^ engineered primary human CD4^+^ and CD8^+^ T cells (total of 6 x 10^6^ T cells) were infused i.v. through tail vein injection. Tumor size was monitored via caliper weekly and tumor weight was measured at completion of experiment. For immunophenotypic analysis, spleens were manually dissociated and subjected to red blood cell lysis (ACK; KD medical).

### LIST OF SUPPLEMENTAL TABLES

**Supplementary Table S1.** ALPPL2 expression in solid tumors. IHC was performed on FFPE tumor tissues using a mouse monoclonal antibody (LifeSpan clone SPM593, catalogue# LS-C390148-20). The number and percentage of ALPPL2 positive cases are indicated. For reference, we also analyzed a public protein expression database, the Human Protein Atlas (https://www.proteinatlas.org), and listed the result of ALPPL2 expression by IHC study using the validated rabbit polyclonal antibody (MilliporeSigma HPA038764). NA: not available.

| Tumor Category | This study | Protein Atlas |
| --- | --- | --- |
| Mesothelioma | 39/91 (42.86%) | N/A |
| Ovarian cancer | 36/60 (60%) | 7/12 (58.33%) |
| Pancreatic cancer | 18/50 (36%) | 2/10 (20%) |
| Gastric cancer | 13/72 (18.06%) | 3/11 (27.27%) |
| Seminoma | 11/11(100%) | 9/11 (81.82%) |

**Supplementary Table S2.**
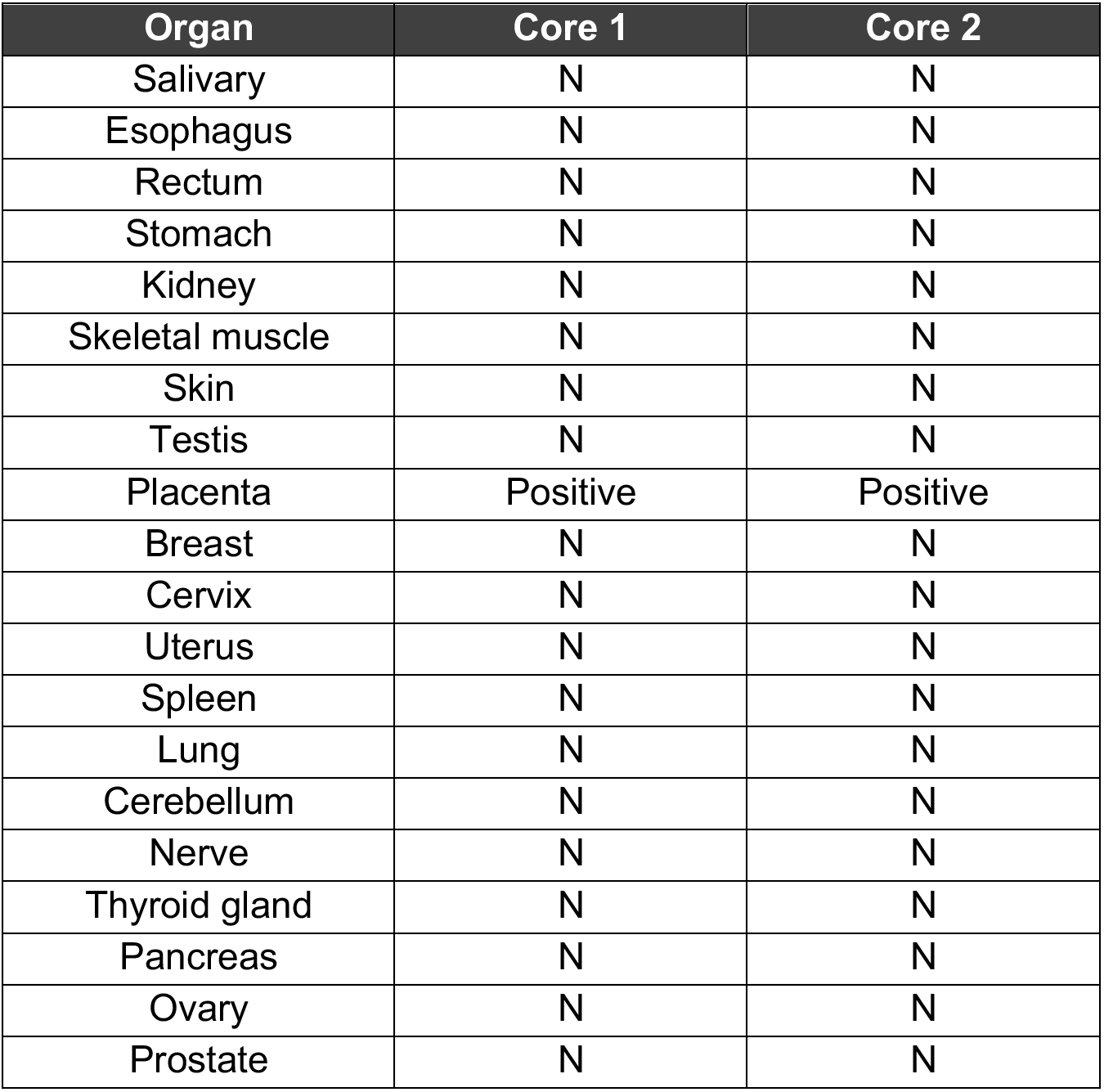
Tissue specificity of ALPPL2 expression. The biotin-labeled M25^FYIA^ scFv and a biotin-labeled non-binding scFv (control) were used to stain normal human frozen tissue arrays. Each tissue type has duplicated cores on the array. Staining results of the M25^FYIA^ scFv were compared with that of the control scFv to determine positive or negative staining. N: negative staining.

**Supplementary Table S3.** IHC study of MCAM expression in FFPE mesothelioma tissue arrays. The number and percentage of each type of staining patterns are indicated. MCAM was detected by both monoclonal (Abcam clone EPR3208, catalogue# ab75769) and polyclonal antibody (pAb) (Abcam, catalogue# ab228487). The difference in percentage between mAb and pAb may reflect difference in sensitivity of antigen/epitope detection in FFPE tissues.

| Staining Result | Anti-MCAM<br>(Rabbit mAb) | Anti-MCAM<br>(Rabbit pAb) |
| --- | --- | --- |
| Positive | 31 (38.27%) | 73 (75.26%) |
| Negative | 50 (61.73%) | 24 (24.74%) |
| Total tissue cores studied | 81 | 97 |

**Supplementary Table S4.**
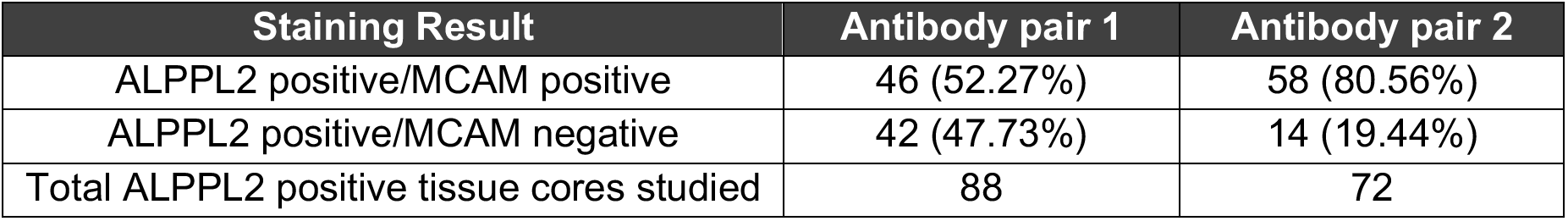
MCAM co-expression in ALPPL2 positive mesothelioma. IHC study using anti-ALPPL2 mouse mAb (LifeSpan clone SPM593) and anti-MCAM rabbit mAb (Abcam clone EPR3208) or pAb (Abcam) was performed on FFPE tumor tissue arrays. Antibody pair 1: Anti-ALPPL2 mouse mAb + Anti-MCAM rabbit mAb. Antibody pair 2: Anti-ALPPL2 mouse mAb + Anti-MCAM rabbit pAb. The number and percentage of each type of staining patterns are indicated.

### SUPPLEMENTAL FIGURES

**Supplementary Figure 1.**
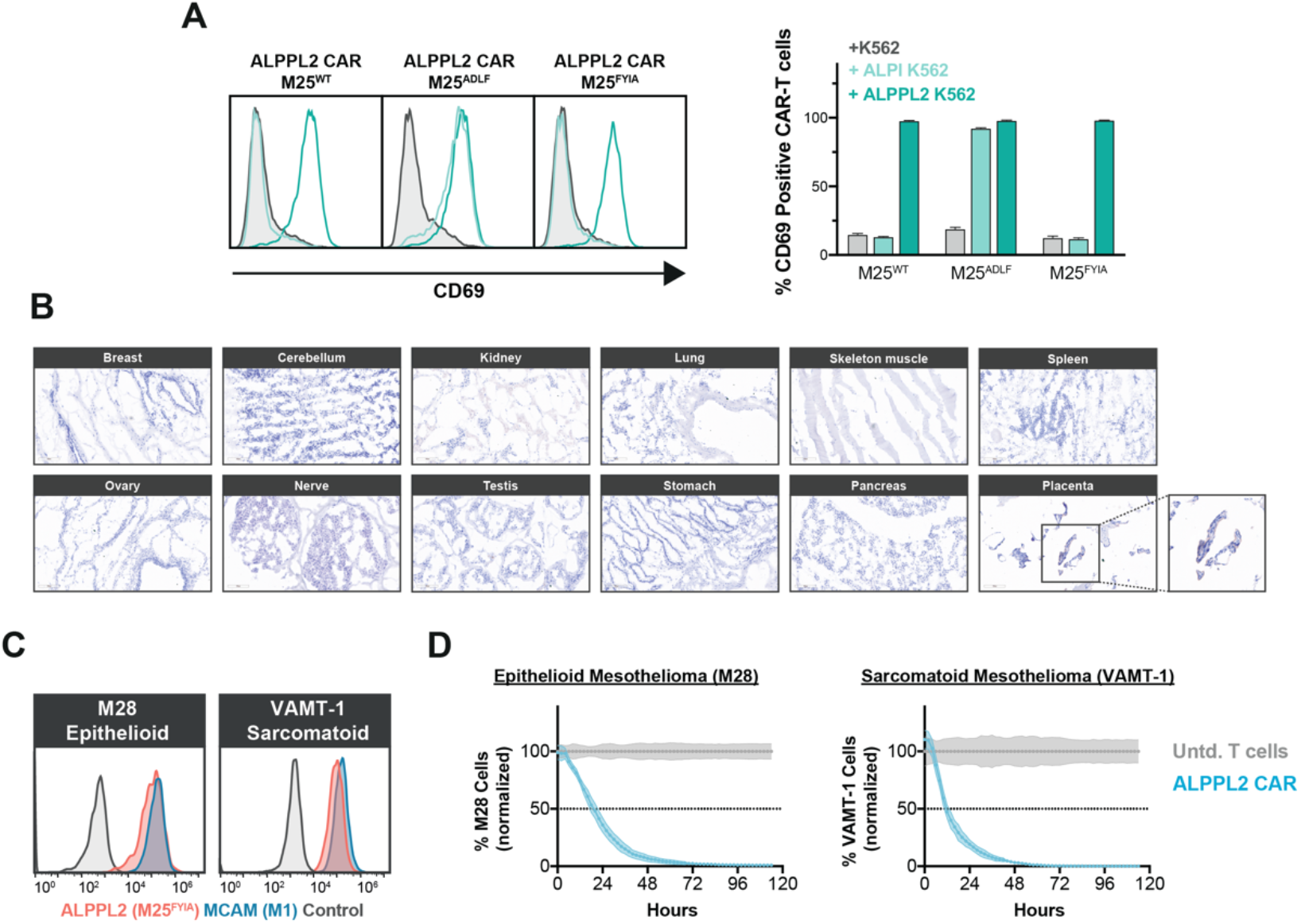
ALPPL2 Tissue Expression and ALPPL2 CAR Specificity. (**A**) Expression of the early activation marker CD69 in CD8^+^ CAR-T cells expressing a BBζ CAR with either a M25^WT^, M25^ADLF^, or M25^FYIA^ scFv 24 hrs after stimulation with K562s expressing either ALPI or ALPPL2. (**B**) IHC study of ALPPL2 expression in normal human frozen tissues using the M25^FYIA^ scFv. Binding was only observed in placental trophoblasts. Scale bar: 100 μm. (**C**) Cell surface co-expression of ALPPL2 and MCAM in an epithelioid mesothelioma cell line M28 and a sarcomatoid cell line VAMT-1 using M1 and M25^FYIA^ scFvs. (**D**) Killing kinetics of epithelioid (M28) and sarcomatoid (VAMT-1) mesothelioma tumor cells by CD8^+^ T cells expressing an ALPPL2 CAR (M25^FYIA^).

**Supplementary Figure 2.**
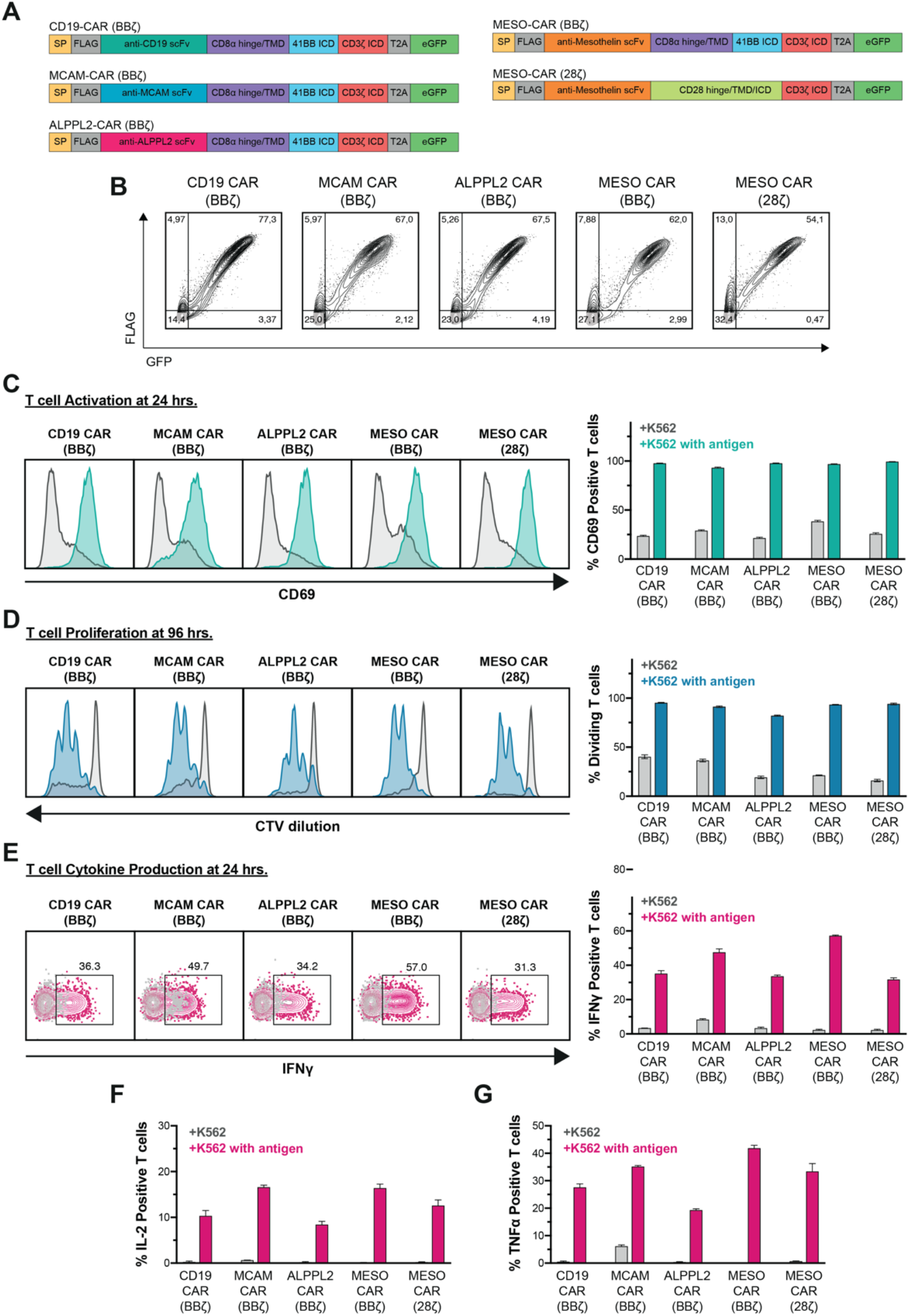
Functional Characterization of Novel CARs. (**A**) Design of CD19, MCAM, ALPPL2, and Mesothelin CARs included in the study and (**B**) their expression in CD8^+^ T cells. SP = signaling peptide; FLAG = FLAG tag; ICD = intracellular domain; TMD = transmembrane domain. (**C**) Expression of the early activation marker CD69 in CD8^+^ CAR-T cells 24 hrs after stimulation with K562s expressing the targeted antigen. (**D**) Proliferation of cell trace violet (CTV) labelled CD8^+^ CAR-T cells four days after stimulation K562s expressing the cognate antigen. (**E**) IFNγ (**F**) IL-2 and (**G**) TNFa production in CD8^+^ CAR-T cells 24 hrs after stimulation with K562s expressing the cognate antigen.

**Supplementary Figure 3.**
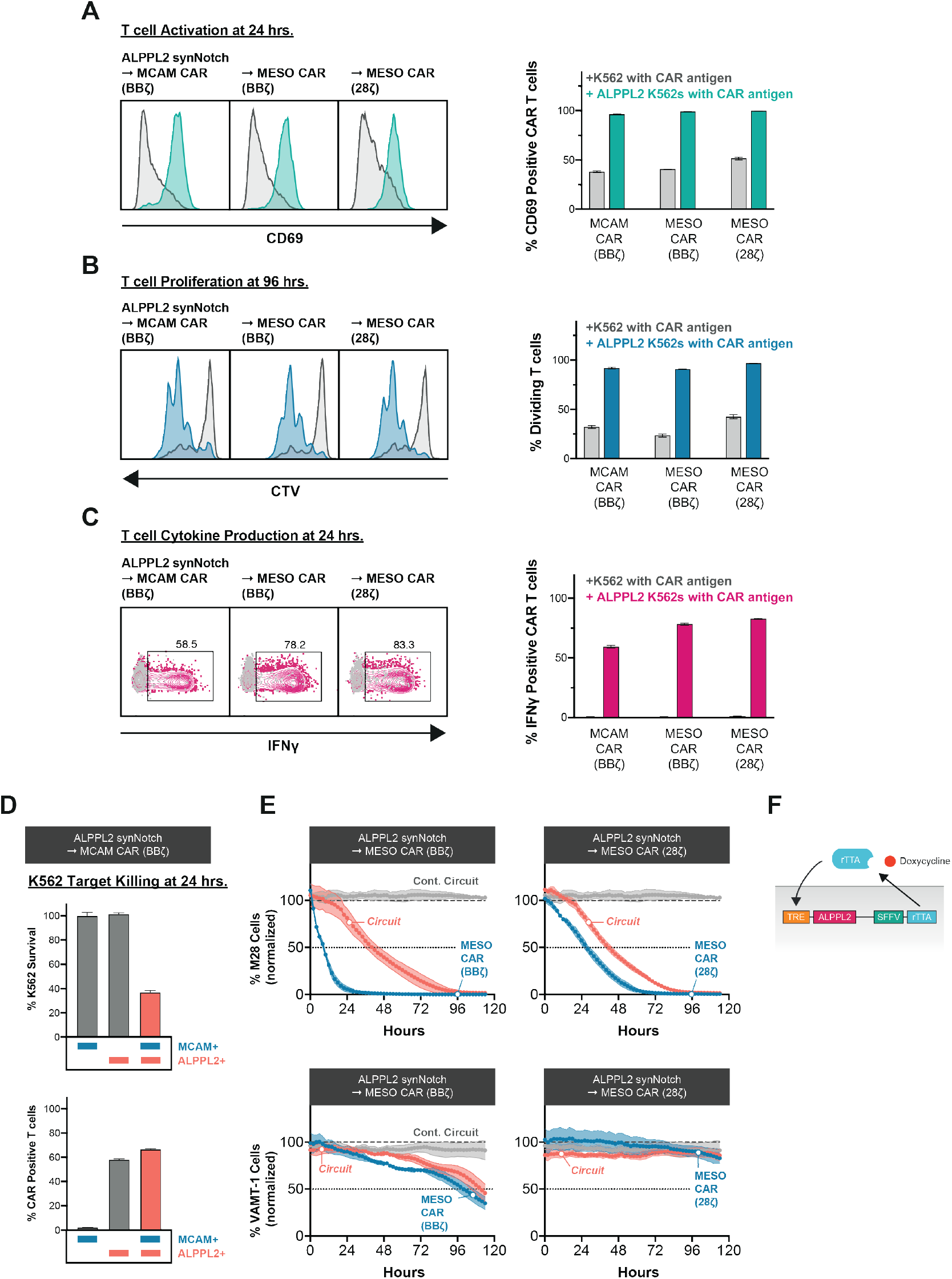
Functional Characterization of ALPPL2-synNotch CAR circuits. (**A**) CD8^+^ T cells were engineered to express an ALPPL2 synNotch together with a genetic circuit encoding for either a MCAM (BBζ) or mesothelin (BBζ or 28ζ) CAR analyzed for CD69 expression in CAR positive cells after 24 hrs of stimulation with K562s expressing the cognate CAR antigen +/− ALPPL2. (**B**) Proliferation of CTV labelled CD8^+^ T cells ALPPL2 synNotch CAR circuits four days after stimulation with K562s expressing the cognate CAR antigen +/− ALPPL2. (**C**) IFNγ production in CD8^+^ T cells ALPPL2-synNotch CAR circuits four after stimulation with K562s expressing the cognate CAR antigen +/− ALPPL2. (**D**) Target killing of K562s expressing MCAM, ALPPL2, or the combination of the two by CD8^+^ T cells expressing an ALPPL2 synNotch MCAM CAR (BBζ) circuit after 24hrs of co-culture. Percentage CAR positive T cells was determined by GFP expression. (**E**) Incucyte assay showing killing kinetics of epithelioid (M28) and sarcomatoid (VAMT-1) mesothelioma tumor cells by CD8^+^ T cells expressing a BBζ mesothelin- or 28ζ mesothelin CARs constitutively or through an. (**F**) Genetic circuit for doxycycline inducible ALPPL2 surface expression.

**Supplementary Figure 4.**
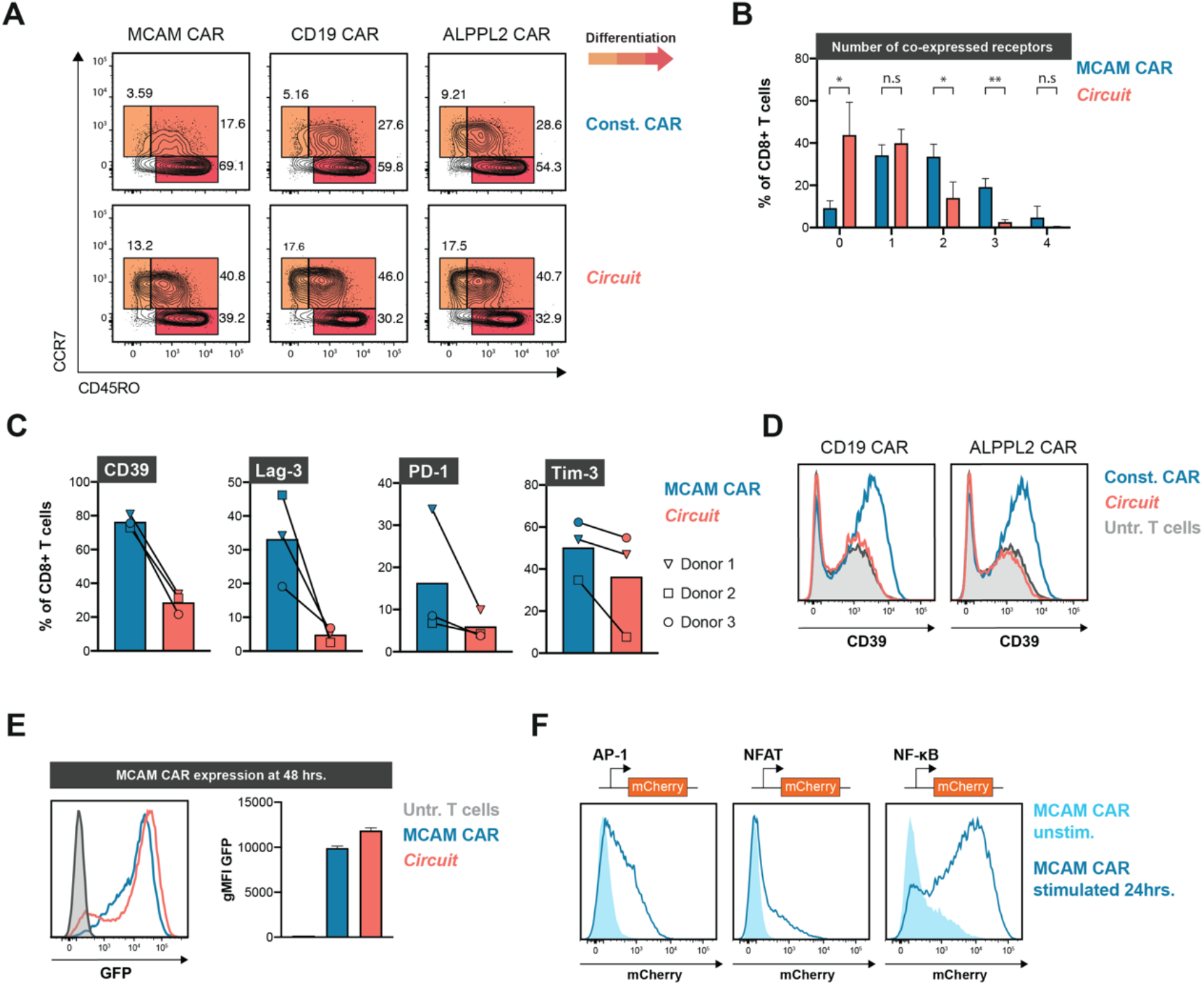
Phenotypic Analysis of CAR and SynNotch CAR T cells Prior to Antigen Exposure. (**A**) Composition of T_SCM_, T_CM_, and T_EM_ in non-antigen exposed CD8^+^ T cells engineered to express a MCAM, CD19, or ALPPL2 CAR either constitutively or through ALPPL2 synNotch circuit 14 days post initial CD3/CD28 Dynabead stimulation. Data linked to figure Fig. 2C. (**B**) Co-expressional pattern and (**C**) fraction of cells positive for CD39, Lag-3, PD-1, Tim-3 in non-antigen exposed CD8^+^ T cells from three different donors engineered to express a MCAM CAR either constitutively or through ALPPL2 synNotch circuit 14 days post initial CD3/CD28 Dynabead stimulation. Data linked to Fig. 2D. (**D**) Expression of CD39 in CD8+ T cells engineered to express a CD19-, ALPPL2 CAR either constitutively or through ALPPL2 synNotch circuit 14 days post initial CD3/CD28 Dynabead stimulation. (**E**) MCAM CAR expression levels in CD8^+^ T cells stimulated with K562s expressing both MCAM and ALPPL2 for 48 hrs as determined by GFP tethered to the CAR. (**F**) AP-1, NFAT, or NF-kB response element activity in Jurkat cells constitutively expressing a MCAM-CAR 24 hrs after stimulation with MCAM positive K562s. Statistics were calculated using unpaired (**B**) or paired (**C**) Student’s t-test. Data is shown as mean±SD. *P≤0.05; **P≤0.01; n.s.; not significant.

**Supplementary Figure 5.**
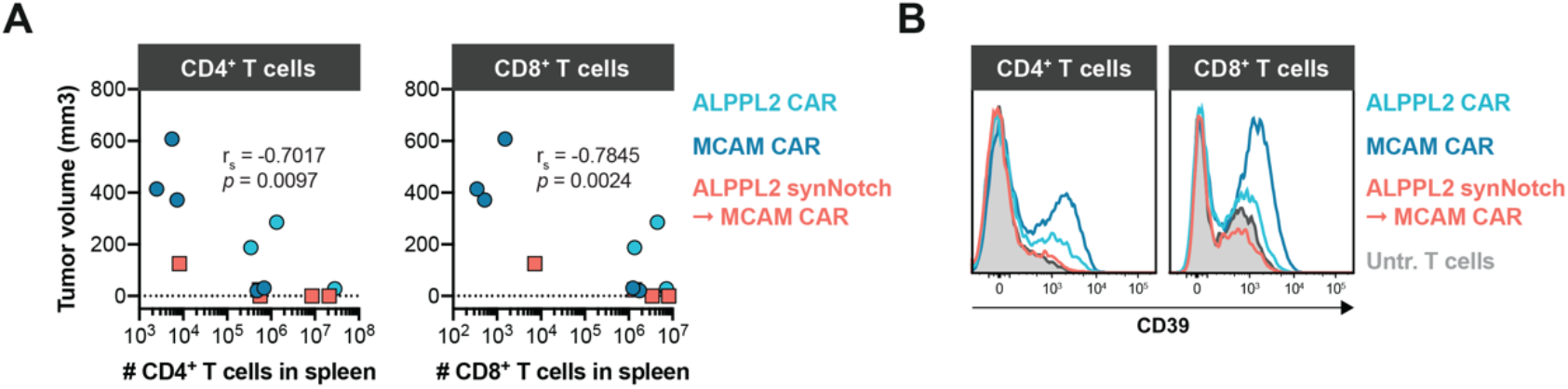
Persistence of CAR T cells and SynNotch CAR Circuit T cells in Treated Tumor-bearing Mice. (**A**) Correlation between final tumor volume and number of human CD4^+^ or CD8^+^ T cells in the spleen at take down. Correlation was determined using Spearman’s rank correlation coefficient. (**B**) CD39 expression in CD4^+^ and CD8^+^ T cells engineered with an ALPPL2 CAR, MCAM CAR, or ALPPL2 synNotch regulating MCAM CAR expression 10 days post initial CD3/CD28 Dynabead stimulation.

